# Family-Based Haplotype Estimation and Allele Dosage Correction for Polyploids Using Short Sequence Reads

**DOI:** 10.1101/318196

**Authors:** Ehsan Motazedi, Richard Finkers, Chris Maliepaard, Dick de Ridder

## Abstract

DNA sequence reads contain information about the genomic variants located on a single chromosome. By extracting and extending this information (using the overlaps of the reads), the haplotypes of an individual can be obtained. Adding parent-offspring relationships to the read information in a population can considerably improve the quality of the haplotypes obtained from short reads, as pedigree information can compensate for spurious overlaps (due to sequencing errors) and insufficient overlaps (due to shallow coverage). This improvement is especially beneficial for polyploid organisms, which have more than two copies of each chromosome and are therefore more difficult to be haplotyped compared to diploids. We develop a novel method, PopPoly, to estimate polyploid haplotypes in an F1-population from short sequence data by considering the transmission of the haplotypes from the parents to the offspring. In addition, PopPoly employs this information to improve genotype dosage estimation and to call missing genotypes in the population. Through realistic simulations, we compare PopPoly to other haplotyping methods and show its better performance in terms of phasing accuracy and the accuracy of phased genotypes. We apply PopPoly to estimate the parental and offspring haplotypes for a tetraploid potato cross with 10 offspring, using Illumina HiSeq sequence data of 9 genomic regions involved in plant maturity and tuberisation.

## 1. Introduction

Genetic polymorphism is the key to understanding inheritance patterns of traits and to identifying genomic regions that affect a trait. Polymorphic genomic loci are used as markers to show co-segregation of genetic variants (alleles) with traits such as resistance to diseases in pedigreed populations, or to find out associations between alleles and the relative abundance of traits in natural populations. These markers can also be used to investigate the genetic components of quantitative (continuous) traits such as height and weight. The sequence of marker alleles along a single chromosome is called a *haplotype*, of which a diploid organism possesses *k* = 2 versions while a polyploid has *k >* 2. To *phase* markers means to determine these *k* haplotypes, which might be identical (harbouring the same alleles) or different (having different alleles at some or all of the marker positions).

Among various types of genetic markers, Single Nucleotide Polymorphism (SNP) markers [1] are the most abundant and are extensively used in genetic studies [2, 3]. While high-throughput assays such as SNP arrays exist for efficient determination of SNP alleles at single loci, direct determination of haplotypes usually requires laborious and expensive techniques such as bacterial cloning, allele-specific PCR or chromosome microdissection [4–6]. However, unphased SNPs provide less knowledge about an individual’s phenotype compared to phased SNPs, as both gene expression and protein function can be affected by an allele being in *cis* or *trans* with other alleles [7]. Moreover, haplotypes can be used as multi-allelic markers offering more statistical power compared to single SNPs for genetic studies [8, 9].

Single individual haplotyping (SIH) methods use DNA-sequence reads to phase the SNPs of a single organism at positions covered by the reads, using the fact that the sequence of called alleles should be the same in the reads that originate from the same chromosome. To deal with sequencing errors, which can cause spurious differences between reads of the same chromosome, these methods use probabilistic models or cost functions to prefer a certain phasing to others based on the observed reads [10–15].

Recently, algorithms have been proposed that apply the rules of Mendelian inheritance to combine the information of reads and pedigree in a cost function for diploids [16] or in a probabilistic model with arbitrary ploidy levels [17]. However, both of these approaches focus on trios, i.e. units consisting of two parents and one offspring, and therefore ignore the information provided by larger population, e.g. in the case of high occurrence of some haplotypes across a large set of progeny which can ease the detection of those haplotypes. In addition, these methods accept recombinant haplotypes in the phasing estimate of the offspring (with the recombination cost/probability being preset as desired), while recombination events are biologically improbable between loci that are only a few thousands nucleotides apart, i.e. in the typical range of haplotypes obtained from short sequence reads. Sequencing and genotype calling errors can therefore be misinterpreted as recombination events by these methods and thus result in spurious haplotypes, especially in polyploids.

Here we propose a new haplotype estimation algorithm, “PopPoly”, that specifically targets larger F1-populations, consisting of two parents and several offspring sequenced by short read sequencing technologies. Considering the short length of the reads, and hence the limitation of read-based phasing to a few hundreds to thousands of nucleotides, PopPoly is based on the assumption that all of the population haplotypes must be present in the parents. Therefore, all of the population reads are combined to estimate the parental haplotypes using a Bayesian probabilistic framework in the first step, and the offspring haplotypes are selected from the estimated parental haplotypes using the minimum error correction (MEC) criterion [18]. In addition, PopPoly uses the pedigree information to detect and correct wrongly estimated SNP dosages and to estimate missing genotypes in the population.

Through extensive simulations, we compare PopPoly to other haplotype estimation methods and show that it improves phasing and variant calling accuracy. Also, we apply PopPoly to estimate haplotypes of plant maturity and tuberisation loci in a cross of tetraploid potato with 10 offspring sequenced with Illumina HiSeq X Ten technology.

## 2. Material and Methods

Short-read sequencing technologies, such as Illumina, produce high-quality sequence reads of up to a few hundred bases in length, which are randomly positioned over the target genomic region and together cover each target position multiple times. By aligning the reads to some consensus reference, genomic variations can be detected and the variant alleles can be specified within each read. To resolve the succession of genomic variants on each chromosome, haplotype estimation or ‘haplotyping’ methods aim to group the reads that have the same variants at the same positions as originating from the same chromosome. This approach requires overlap of the reads at the variation sites and the inclusion of at least *two* variation sites in a read, so that the flanking positions can be connected by the overlaps at the position(s) in between.

However, some of the reads do not meet the criterion of containing at least two variation sites, and the connection between the variation sites can be therefore broken at some positions. For this reason, current haplotyping algorithms start by detecting positions connected to each other through the sequence reads and aim to resolve the haplotypes over each obtained set of connected positions, i.e. the so-called "haplotype blocks" or solvable islands. With short sequence reads, haplotype blocks often include a few hundred up to a few thousand bases.

In our approach, we use the fact that recombination events are usually extremely unlikely over the short distances covered by the haplotype blocks obtained from short reads. Therefore we combine all of the reads in an F1-population to estimate the parental haplotypes, and determine the haplotypes of each offspring by selecting the phasing the most compatible with its reads from the set of phasings possible by the transmission of the (already estimated) parental haplotypes.

To implement this method, we follow a greedy SNP-by-SNP extension approach (Figure 1), extending the base phasings *H_bm_* and *H_bf_* (for the mother and father, respectively) at each step by one SNP and choosing the most likely phasing extensions *H_em_* and *H_ef_* to continue with as the base phasings of the next step until all of the *l* SNPs within a haplotype block have been phased. Starting by the first two SNP positions in the block, the probabilities of the base and extended phasings, conditional on the reads and taking offspring genotypes into account, are calculated using the Bayes formula. Finally, the offspring haplotypes are chosen from the estimated parents using the minimum error correction (MEC) criterion, so that the phasing selected for each offspring have the maximal compatibility with its individual reads (Section 2.1). A natural advantage of such an approach is that the uncalled SNP genotypes of an offspring are imputed in its haplotypes if those SNPs are included in the parental phasings. In addition, the Bayesian framework for phasing extension can be used to detect erroneous SNP genotypes, which result in zero probabilities for all extensions at a SNP position. We use a similar Bayesian approach to re-estimate these erroneous genotypes, as well as the uncalled SNP genotypes of the parents, by assigning probabilities to the possible genotypes at a SNP position conditional on the reads and the parent-offspring relationships from which the most likely genotype is chosen as the estimate (Section 2.2).

**Figure 1.**
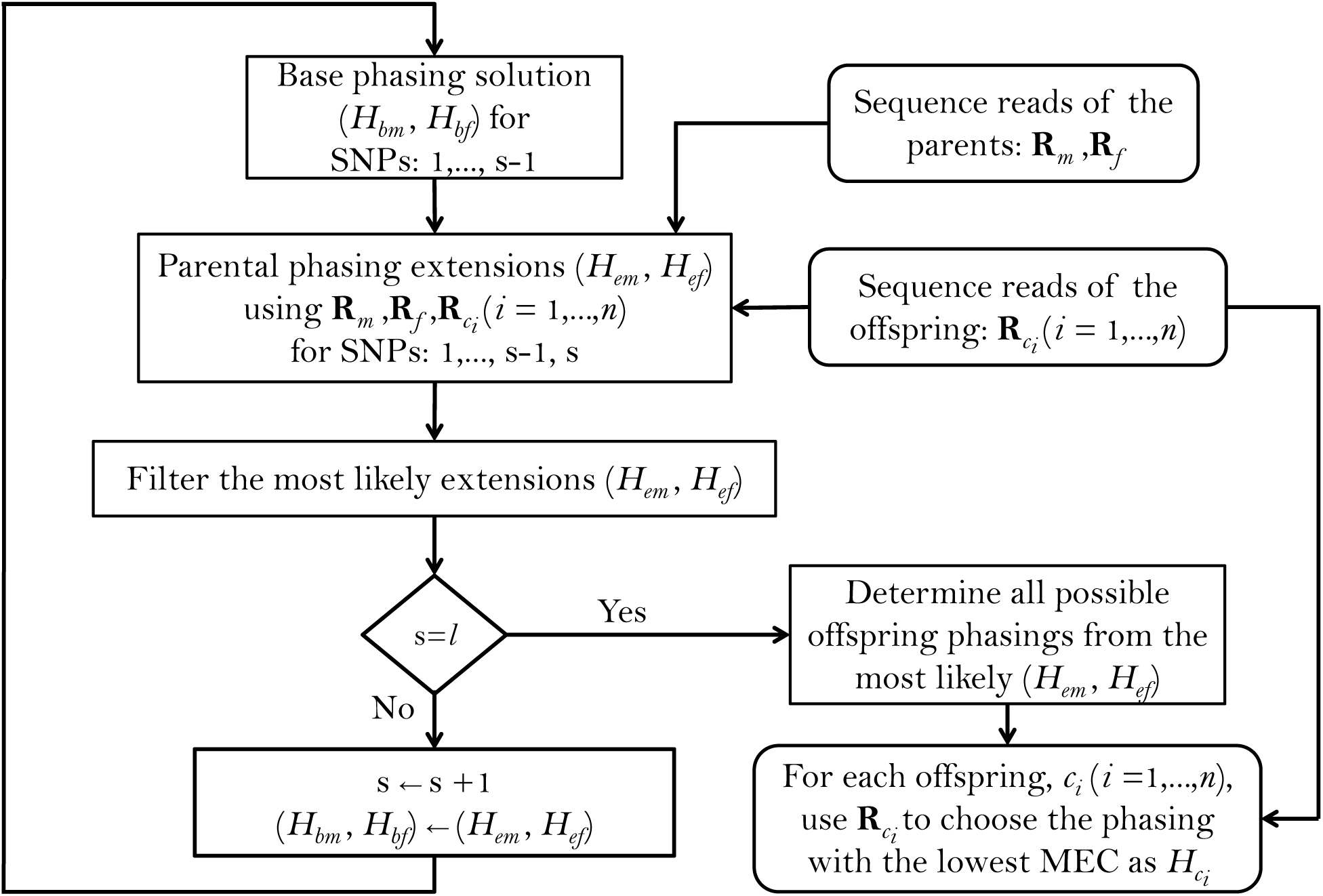
Summary of the “PopPoly” method to estimate haplotypes in an F1-population with two parents, (*m*, *f*), and *n* offspring, *c_i_* (*i* = 1*,…, n*), using the sequence reads for a connected region including *l* SNPs.

### 2.1. Estimation of parental haplotypes

Inspired by the approach of Berger *et al*. [12], we start at the first SNP position in the target region (*s* = 1), and extend the maternal and paternal genotypes of this SNP, 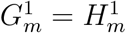 and 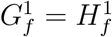, respectively, to two-SNP phasings, 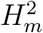 and 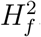. We consider every possible phasing between 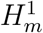 and 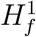 and SNP position *s* = 2 in the region, and obtain the joint conditional probability of each extension pair, (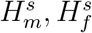), at *s* = 2 given the sequence reads of the population and the parental genotypes, (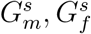), as well as the offspring genotypes 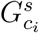 for *i* = 1*,…,n* (with *n* representing the number of offspring). Keeping only those parental extensions whose conditional probability exceeds or equals a pre-set *branching* threshold, *ρ* ∈ (0, 1], we eliminate further the extensions whose probability is less than *κP_max_*, where *κ* ∈ [0, 1] is a preset *pruning* threshold and *P_max_* is the maximum probability assigned to the candidate parental extensions. The surviving extensions at *s* = 2 are used in the next step as base phasings to obtain the extensions at *s* = 3 in a similar manner, and this procedure is continued until the last SNP *s* = *l* has been added to the parental extensions.

As it is not straightforward to directly calculate the conditional extension probabilities [17], we calculate instead the probability of the sequence reads conditional on each possible phasing and convert these probabilities to the desired extension probabilities using Bayes’ formula:

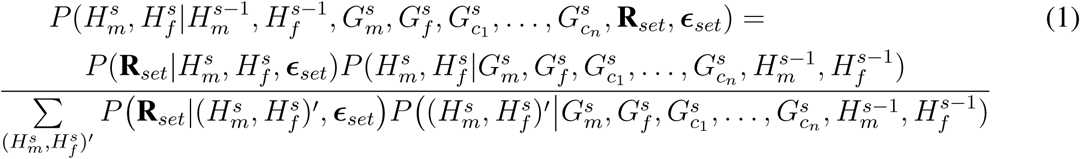

where **R**_*set*_denotes the set of all of the reads in the population and ***ϵ***_*set*_stands for the set of base-calling error vectors, *ϵ_j_*, associated with each *r_j_* ∈ **R**_*set*_(1 ⩽ *j* ⩽ *|***R***_set_|*). 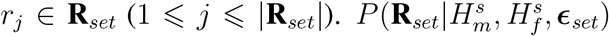 denotes the conditional probability of observing the reads given a pair of maternal and paternal extensions at *s*, 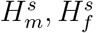, and the base-calling error probabilities given by ***ϵ***_*set*_.

To calculate 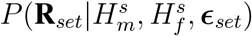, we assume conditional independence of each read, *r_j_* ∈ **R**_*set*_, from the other reads in **R**_*set*_given ***ϵ***_*set*_, and use the fact that each read is either directly obtained from one of the parental samples or belongs to an offspring *c_i_* (*i* = 1*,…, n*), in which latter case the read may have originated from either parent with equal probability. Under these assumptions, 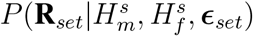 is determined according to:

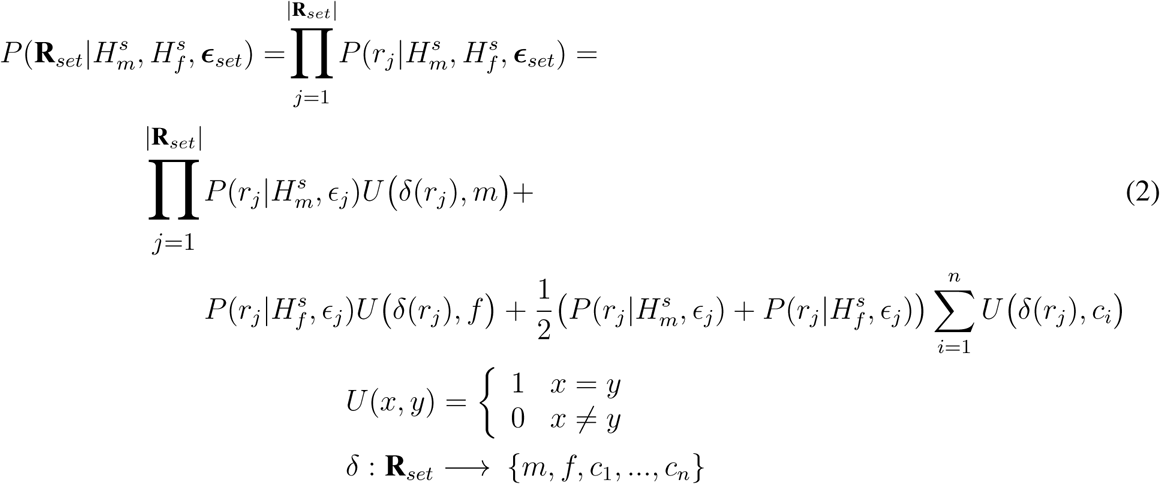

where the function *δ*(*r_j_*) returns the origin of read *r_j_*: mother (*m*), father (*f*), or one of the *n* offspring (*c*_1_*,…*.*, c_n_*).

Assuming independence of the sequencing errors at the SNP positions within each read, 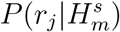 and 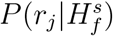 in Equation 2 can be calculated according to [17]:

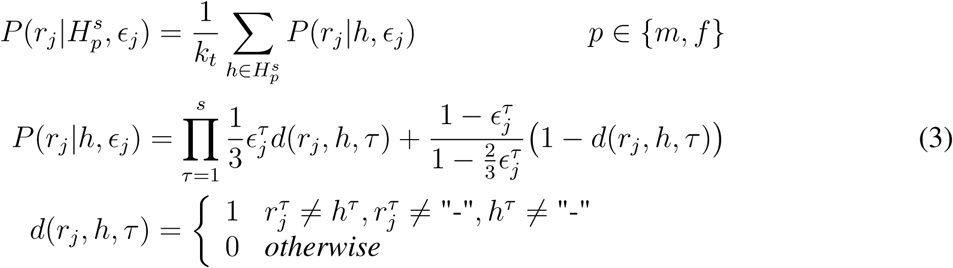

where *ϵ_j_* assigns a base-calling error probability to every SNP position in *r_j_*, and *h* stands for each of the *k_t_* homologues in the phasing extension 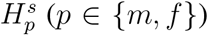. In Equation 3, we use the superscript *τ* in 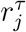 and 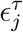 to represent the called base at SNP position *τ* and its associated error probability, respectively. Likewise, *h^τ^* denotes the allele assigned to homologue *h* at SNP position *τ*. We use 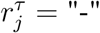 and *h^τ^* = ‘-’ to show that SNP position *τ* has not been called in *r_j_* or is missing in *h*.

Equations 2 and 3 establish the procedure to calculate the likelihood in Bayes’ formula in Equation 1. In order to solve Equation 1, one also needs to specify the prior, 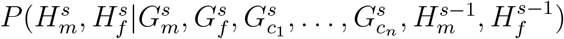. While several ways can be thought of to specify this prior, we obtain it as follows.

As the parental extensions (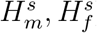) are confined to those compatible with 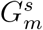 and 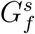, we set this prior to zero for every incompatible extension. For the compatible extensions, we look into the possible transmissions of the extended haplotypes (ignoring phenomena like aneuploidy [19], preferential chromosome pairing [20], recombination and double reduction [21]) to the offspring and for each offspring, *c_i_*, we count the number of transmissions that agree with its genotype at *s*, 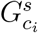. Dividing this number by the total number of possible transmissions, (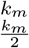) · (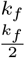), gives us 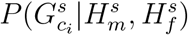. Calculating 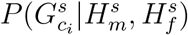 for *i* = 1,…,*n*, we obtain the average likelihood of an *observed* offspring genotype according to:

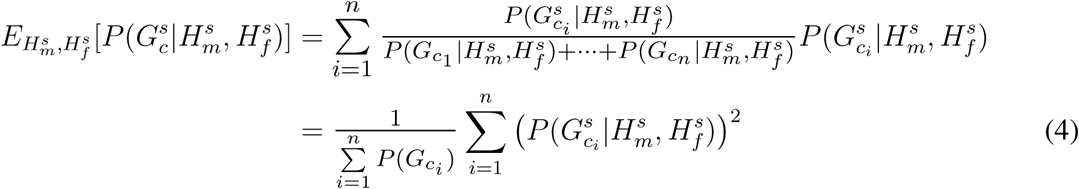

where 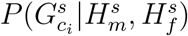 is the likelihood and 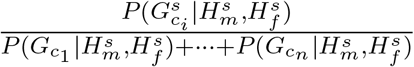 is the probability of observing offspring *c_i_*.

So far, we set the prior for each (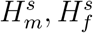) to be proportional to 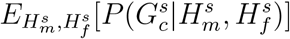. However, as changing the order of the homologues does not change a phasing, several permutations of the alleles at *s −* 1 and *s* can yield the same (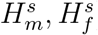). Therefore, the prior should also be proportional to the number of permutations that result in (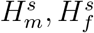). It can be thus set to:

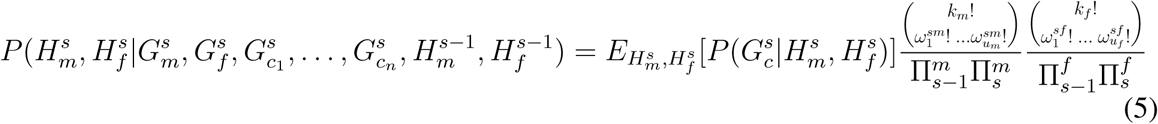

where, for 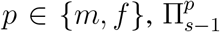 and 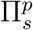 are the number of possible permutations of the alleles at *s−*1 and *s*, respectively, *u_p_* is the number of distinct homologues, i.e. haplotypes, in 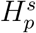 regarding only positions *s –* 1 and *s*, and 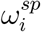 for *i* ∈ {1*,…, u_p_*} denotes the number of times an identical haplotype (regarding only positions *s* − 1 and *s*) is present in 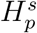. Although it is possible to normalise the priors obtained this way over all of the possible extensions (to obtain a proper prior mass function), one does not need to do so as the discrete posteriors are normalised anyway at the end.

As an example, with tetraploid parents there will be 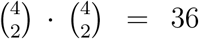 possible haplotype transmissions to each offspring. With maternal and paternal extensions at *s* = 3 being equal to 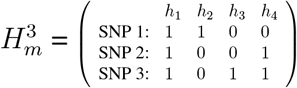 and 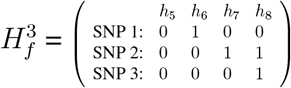, respectively, and two offspring *c*_1_ and *c*_2_ with 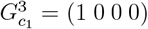 and 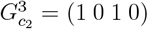, only 9 out of 36 transmissions will be compatible with the genotype of *c*_1_, while 18 transmissions will be compatible with *c*_2_. This results in 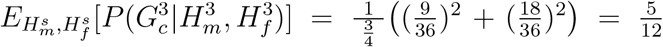 for this extension. As 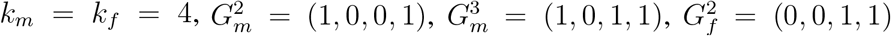 and 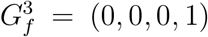 we have 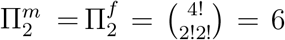 and 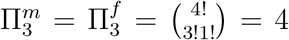. Considering only SNPs at *s −* 1 = 2 and *s* = 3, in each parent there is one haplotype present twice. The a priori probability of (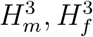) is hence determined from Equation 5 to be 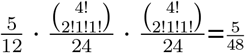.

From Equations 2 and 5, the conditional probabilities of parental extensions at position *s* can be obtained using Equation 1 and the surviving extensions are used for the extension to *s* + 1, as explained above.

### 2.2. Estimation of missing and erroneous genotypes

The SNP-by-SNP extension of the parental haplotypes using the sequencing reads of an F1-population was explained in the previous section, assuming the SNPs have been accurately called for all of the population members. However, in practice every haplotyping algorithm has to handle missing and wrongly estimated SNP genotypes caused by sequencing and variant calling errors.

In presence of wrongly estimated genotypes (wrong dosages), it can occur that all of the offspring genotypes are incompatible with the parental extensions at some SNP position *s*. At these positions, the extension should either be skipped, as the prior weight of all candidate phasings will be zero, or the genotypes must be estimated anew. The extension at *s* will also be impossible if one or both of the parental genotypes are missing at *s*. To include these SNP positions in the extension, it is necessary to impute the missing genotypes.

In order to estimate the population genotypes at the missing or incompatible positions, we assume that the parents come from an infinite-size population at Hardy-Weinberg equilibrium. Limiting the attention to bi-allelic SNPs, the reference and alternative allele frequencies of the parents at position *s* can be estimated from the observed reads under the above assumption. Assuming a fixed sequencing error rate for all of the reads and nucleotide positions, 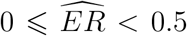, the frequency of the alternative allele can be obtained assuming a binomial model for the observed count of the alternative allele according to:

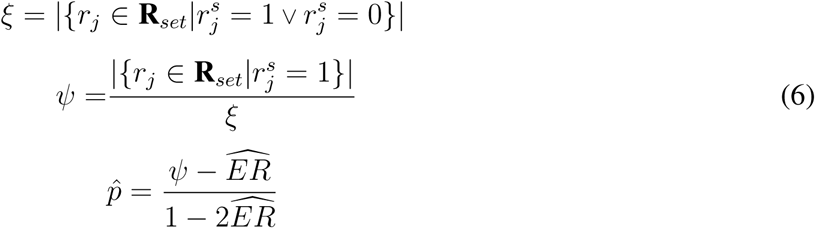

where *ξ* is the total sequencing coverage of the population at *s* and *ψ* is the proportion of the alternative allele among the observed alleles. As this observed frequency, *ψ*, depends on the latent true frequency, *p*̂, through 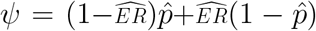, it is straightforward to show that *ψ* can be obtained as shown in Equation 6, with a standard error equal to 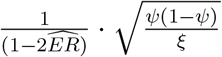.

In case a specific base-calling error rate 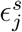 is assigned at each position *s* to each read *r_j_* e.g. by using the integer-rounded Phred (quality) scores reported by the sequencer [22], one can assume a Gaussian distribution for the probability of observing the alternative allele at *s* in each read, 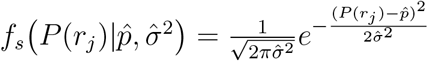, and obtain *p*̂ at each *s* according to:

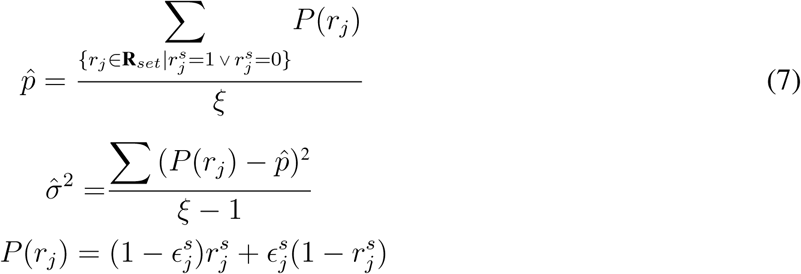

Having *p̂*, a prior probability can be assigned to each of the 2^*k_m_*^ and 2^*k_f_*^ theoretically possible genotypes for the mother and the father, respectively, assuming a binomial model according to:

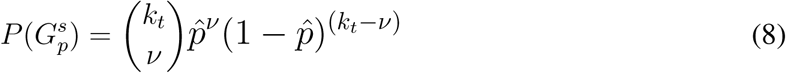

where *p*∈ {*m, f*} and 0 ⩽ *ν* ⩽ *k_t_* is the dosage of the alternative allele in the candidate genotype, 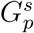. Assuming the parents have been independently chosen from a source population, a prior can be assigned to each (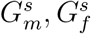) pair using 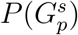 obtained from Equation 8, according to:

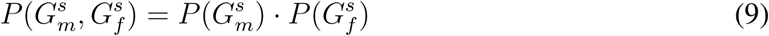

Given (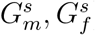), a prior probability can be assigned to each specific offspring genotype, 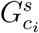, by counting the number of allele transmissions that result in that 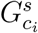. For example, with 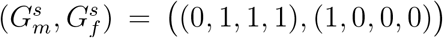 the prior 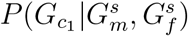 will be equal to 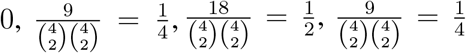 and 0 for the offspring genotypes: *G_c_*_1_ = (0; 0; 0; 0),*G_c_*_1_ = (1, 1, 0, 0), *G_c_*_1_ = (1, 1, 1, 0) and *G_c_*_1_ = (1, 1, 1, 1), respectively.

To estimate the population genotypes, 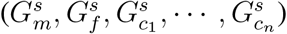, we use the prior probabilities obtained as explained above, and assign a posterior probability to each population genotype by taking the sequencing reads into account. Noting that:

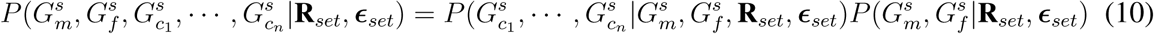

we separately obtain the posterior of the parental genotypes, 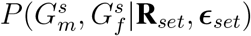, and the conditional posterior of the offspring 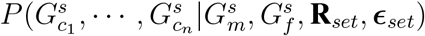, from which the population posterior is derived using Equation 10.

The posterior of (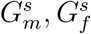) can be directly obtained from Equations 1 and 2 by substituting (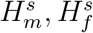) with (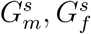) in these equations. Assuming conditional independence of the offspring genotypes given the parents, we obtain 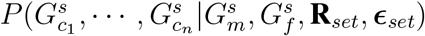 by:

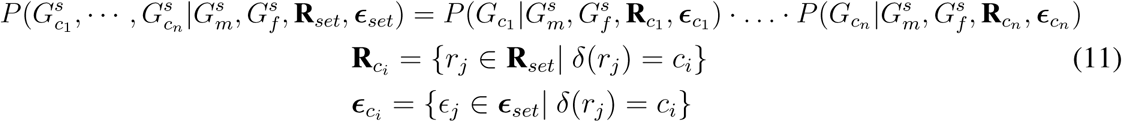

where 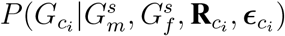 is calculated according to:

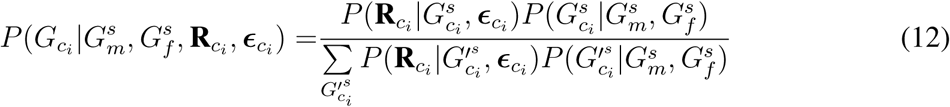

and:

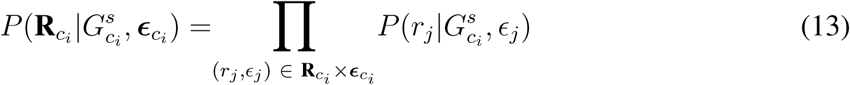

where 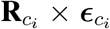 represents the Cartesian product of 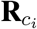 and 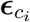, and (*r_j_, ϵ_j_*) denotes 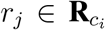 with its matched error rate vector, 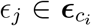. In Equation 13, 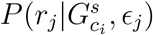 is obtained by replacing 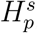 with 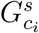 in Equation 3.

After calculating 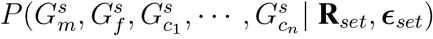 from Equation 10, the most likely population genotypes at *s* can be assigned to the population members as genotype estimates.

### 2.3. Estimation of the offspring haplotypes

Having the set of all possible offspring phasings obtained by the possible transmissions of the parental haplotypes (Section 2.1), we assign to each offspring *c_i_* the phasing estimate 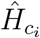 that yields the smallest number of required base-calling changes in the sequence reads, 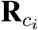, in order to assign each 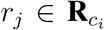 to some homologue in 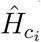. For each possible offspring phasing, *H*̂, this required number of base-calling changes equals the so-called *minimum error correction* (*MEC*) score, defined as [18]:

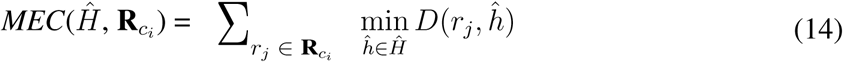

*D*(*r_j_, h*̂) is the Hamming distance between read 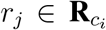 and homologue *h*̂ ∈ *H*̂ defined according to:

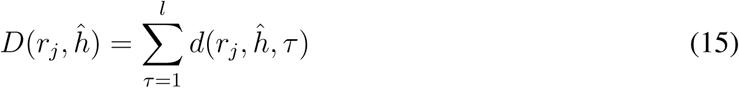

where *τ* and *l* represent the SNP positions and the number of SNPs in the target region, respectively, and *d*(*r_j_, h*̂, *τ*) is defined in Equation 3. Thus, for each *c_i_* we have 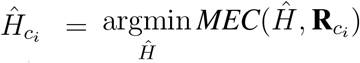. If 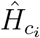 is the same as the true phasing of *c_i_*, its MEC score is expected to be close to the number of actual base-call errors in 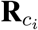.

In case more than one set of parental haplotypes has the maximum probability (Section 2.1), we infer the offspring haplotypes for each of them as explained above and finally choose the family whose offspring MEC score is the smallest.

### 2.4. Performance evaluation by simulation

To evaluate the performance of PopPoly and compare it to other haplotyping methods, we simulated genomic regions of length 1 kb for F1-populations of tetraploid potato, as described in Motazedi *et al*. (2017) [17], introducing on average one SNP per 50 bp in each parental sequence. For the potato genome, typical genetic distances have been reported to be in the range of 3 to 8 *cM/Mb* [23] [21]. Therefore, the assumption of improbable recombination holds for the simulated genomic regions.

We simulated different scenarios varying the number of offspring from 1 to 30, and for each scenario generated *in silico* paired-end Illumina HiSeq 2000 reads, with an average insert-size of 350 bp and single read length of 125 bp, using the sequencing simulator ART [24]. The simulated sequencing depth was 5 × per homologue for each parent and 2 × per homologue for the offspring. We also conducted simulations of families with 2, 6 and 10 offspring with higher sequencing depths, up to 30 × per homologue for each individual, in order to evaluate the performance at higher coverages.

After mapping the simulated reads to their reference regions using BWA-MEM [25] and calling SNPs using FreeBayes [26], we estimated the phasing of the parents and the offspring in each F1-population using SIH methods: SDhaP [13] and H-PoP [15] (shown to perform better than other SIH methods such as HapCompass [11], HapTree [12] and SDhaP), as well as the trio based method available for polyploids: TriPoly [17].

We used several measures to compare the accuracy of haplotype estimation with the used methods. These include the *pair-wise phasing accuracy rate* (*PAR*), defined as the proportion of correctly estimated phasings for SNP-pairs [27], as well as the *reconstruction rate* (*RR*) defined to measure the similarity between the original haplotypes and their estimates at each SNP site [17].

As the quality of haplotype estimation depends not only on the accuracy of the estimated haplotypes, but also on the ability of haplotyping method to phase as many SNPs as possible and to efficiently handle missing SNPs and wrong dosages, we used *SNP missing rate* (*SMR*) and *incorrect dosage rate* (*IDR*) in the estimated haplotypes to get insight about these aspects for each method. Finally, to show the continuity of phasing we measured the average number of phasing interruptions, i.e. the number of haplotype blocks minus one, in the estimates of each method and normalised it by the number of SNPs, *l*, as *number of gaps per SNP* (*NGPS*).

### 2.5. Haplotype estimation of tuberisation and maturity loci in potato

We used PopPoly to estimate haplotypes of the tuberisation and maturity loci reported by Kloosterman *et al.* [28], in an F1-population with 10 offspring obtained from the cross of two *S. tuberosum* cultivars: Altus × Colomba (*A* × *C*). The nine investigated loci (Table 1) belong mainly to the potato cycling DOF factor (*StCDF*) gene family, but also include other genes, such as CONSTANS (CO) genes CO1 and CO2, that are shown to be involved in *StCDF* regulation [28].

**Table 1.**
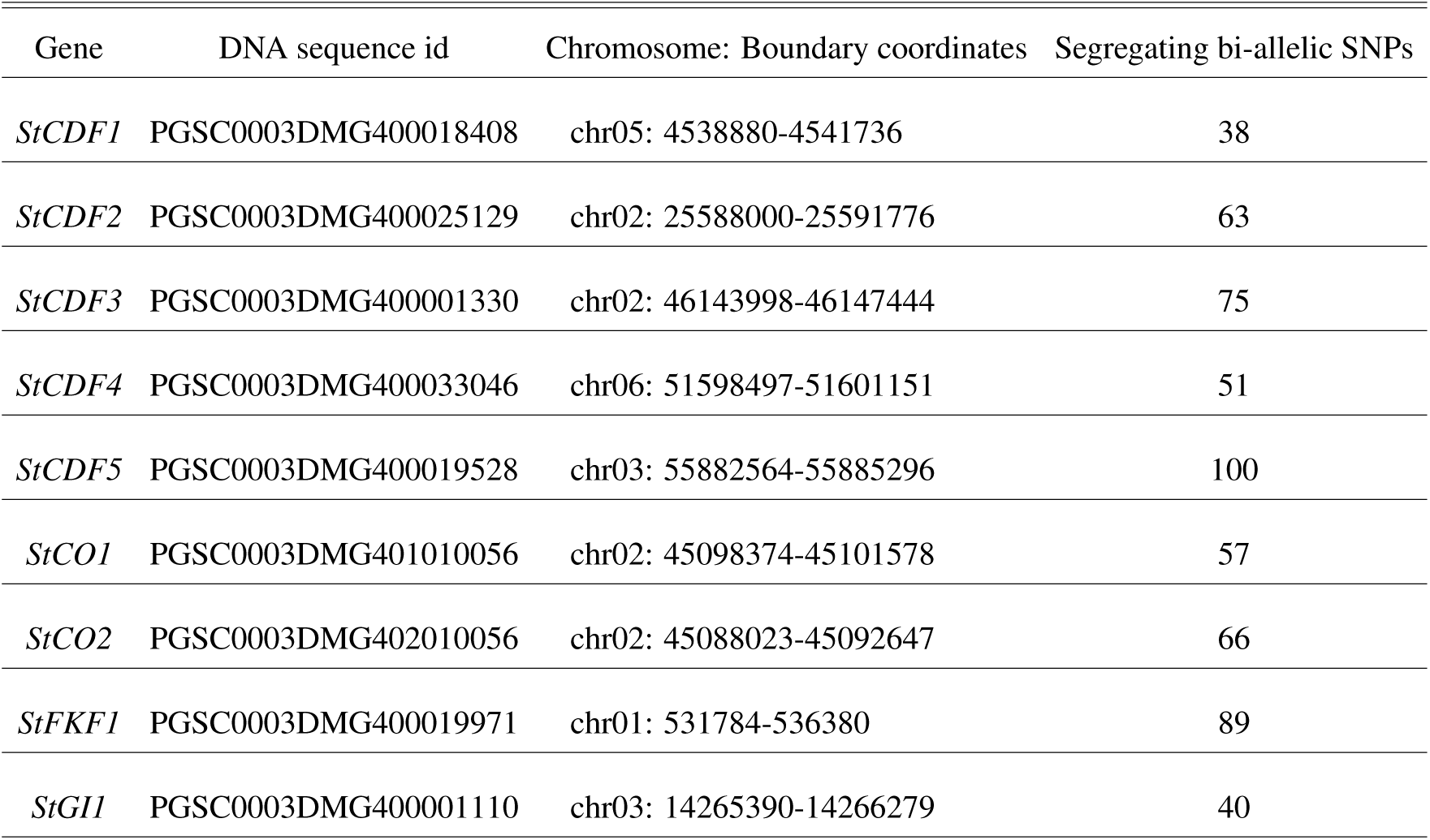
List of the *S. tuberosum* genes selected for haplotyping

Sequence data for the *A × C* population was obtained by whole genome sequencing (WGS) using Illumina HiSeq X Ten technology. Paired-end sequences were obtained with an average insert size of 380 bp (single read length of 151 bp) and aligned to PGSC-DM-v4.03 reference genome [29] using BWA-MEM [25]. Genomic variation within the boundaries of the selected genes was detected from the aligned reads using FreeBayes [26], with an average read depth of 85× (sd=30×) at the target loci. The paired-end sequence reads were used by PopPoly to estimate the phasing of the detected bi-allelic SNP sites (including SNPs obtained by collapsing FreeBayes complex variants).

## 3. Results and Discussion

### 3.1. Simulation study

To evaluate the performance of PopPoly, we simulated potato F1-populations with 1 to 30 offspring and estimated the population haplotypes using PopPoly as well as SDhaP, H-PoP and TriPoly. The estimated haplotypes were compared to the original haplotypes by *hapcompare* [27], using the measures introduced in Section 2.4. The results are summarised below.

#### PopPoly yields more accurate offspring haplotypes

The comparison of the reconstruction rates (RR) reported for the phasing estimates of the offspring showed that RR, which is a measure of overall phasing accuracy, is around 4% higher for PopPoly compared to the to the next most accurate method, TriPoly (Figure 2-a). The second measure of accuracy, the pairwise-phasing accuracy rate (PAR) which is especially sensitive to the accuracy of phasing between distant SNPs, was around 12% higher for the offspring estimates obtained by PopPoly (Figure 2-b) compared to the next method (TriPoly). Together, these two measures show that PopPoly improves the accuracy of phasing in the offspring compared to the other methods.

**Figure 2.**
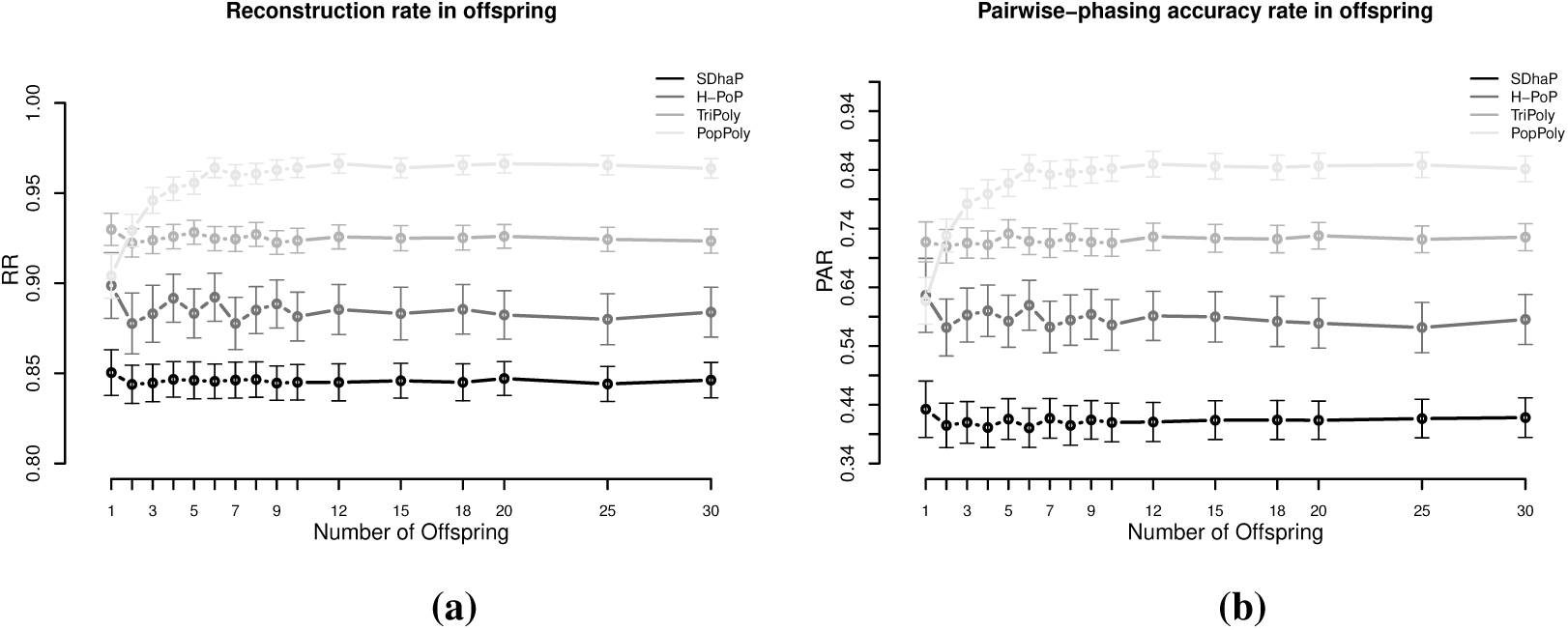
Haplotyping accuracy measures: (a) RR, (b) PAR in the offspring against the number of offspring in the population using PopPoly (light grey), TriPoly (grey), H-PoP (dark grey) and SDhaP (black) for simulated tetraploid potato populations.

However, the accuracy of PopPoly depends on the population size, especially for distant phasing evaluated by PAR. As seen in Figure 2-b, PAR increases rapidly for PopPoly with an increase in the number of offspring from 1 to 3. In fact, the highest offspring score for a trio, i.e. with only one offspring, is reported by TriPoly and the accuracy of PopPoly reaches that of TriPoly when there are at least 2 offspring in the population.

However, the dependence of PopPoly accuracy on the population size gradually diminishes as the number of offspring reaches 6. As an increase in the count of a haplotype in the population results in an increase in the number of reads compatible with that haplotype (assuming no sequencing bias), the power of PopPoly algorithm increases to detect that haplotype. With a trio, however, there is a chance that some of the parental haplotypes are not transmitted to the offspring from a tetraploid cross. This causes the lower accuracy of PopPoly compared to TriPoly when applied to a trio, as the latter method does not combine the offspring reads with the reads from the parents.

While increasing the per homologue coverage from 5-5-2× (mother-father-offspring) to 30-30-30× yielded an average increase of 23-36% in PAR for TriPoly, H-PoP and SDhaP, the increase was only 14% for PopPoly (Supplement S3), as combining the population reads effectively augments the haplotyping coverage (the increase was actually less than 5% with 10 offspring, Supplement S3). Similarly, the difference in RR between the lowest and the highest coverage was 3% for PopPoly compared to 4-6% for the other methods.

For the parents, the reported accuracy measures were very similar between the methods, with H-PoP and PopPoly yielding the highest scores (Supplement S1).

#### Haplotype estimates of PopPoly include more SNPs compared to the other methods

As shown in Figure 3, the average SNP missing rates (SMR) of PopPoly are around 20% lower compared to H-PoP and around 9% lower compared to TriPoly and SDhaP. The reason for this is that combining individual NGS reads increases the chance to phase parental SNPs and choosing the offspring phasings from the estimated parental haplotypes leads to the inclusion of SNPs not sufficiently covered by the offspring reads, as well as to the imputation of SNPs uncalled in (some of) the offspring.

**Figure 3.**
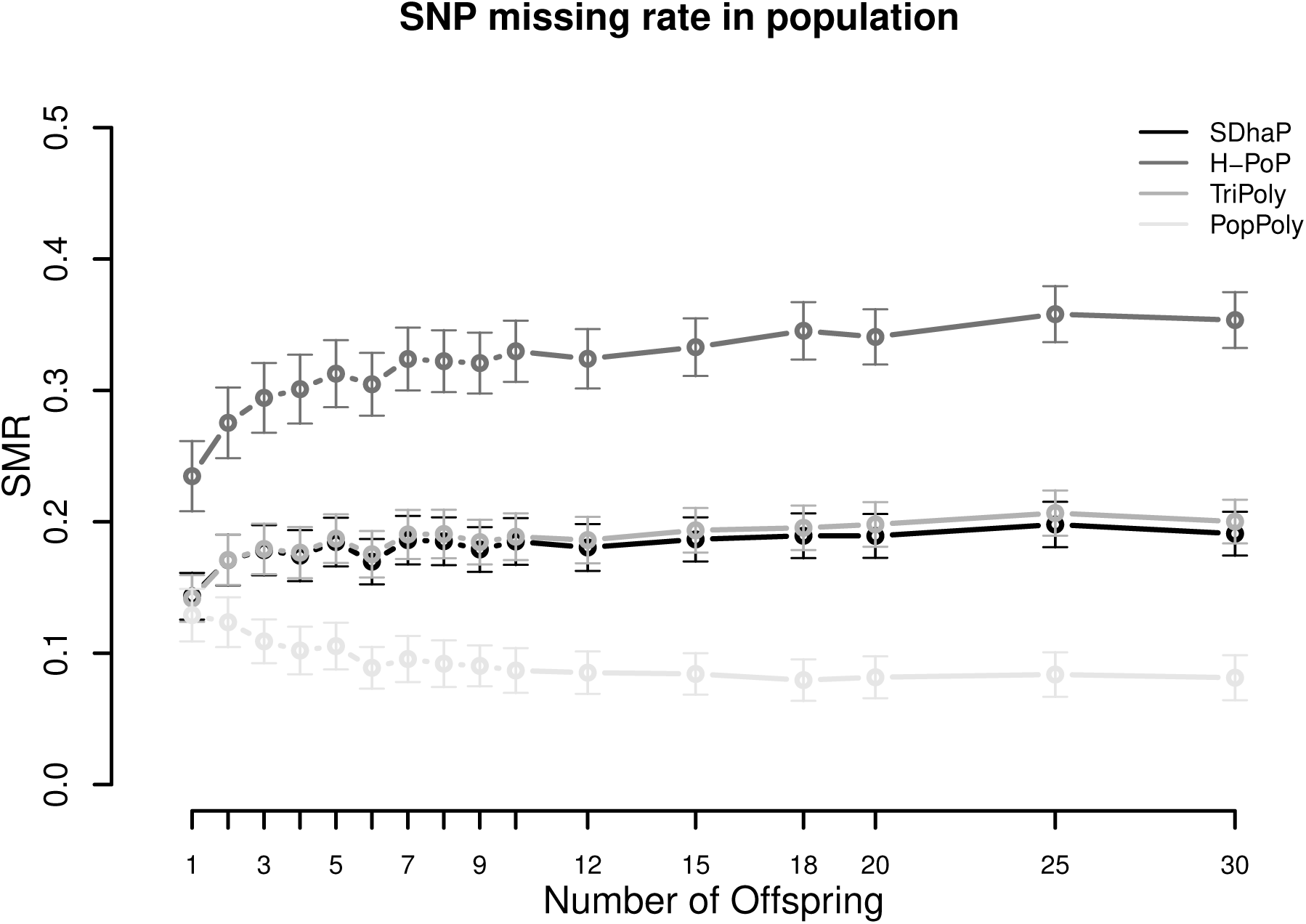
SNP missing rate (SMR) in the population against the number of offspring reported by PopPoly (light grey), TriPoly (grey), H-PoP (dark grey) and SDhaP (black) for simulated tetraploid potato populations.

However, around 10% of SNPs are still missing in the PopPoly phasing, as PopPoly excludes a SNP position if the offspring genotypes at that position (either given as input or estimated anew by PopPoly) are incompatible with the surviving parental extensions. An example of this for a trio is the extension at *s* = 2, if the only surviving parental extensions are 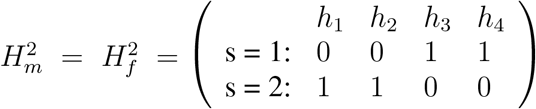 and the offspring genotype at *s* = 2 is 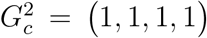. While 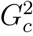 is compatible with the parental genotypes at *s* = 2 (and therefore is accepted by the point-wise dosage estimation of PopPoly), it cannot be obtained from 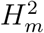 and 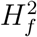 without meiotic recombination. Since PopPoly is based on the assumption of no recombination (Section 2.1), it excludes this SNP site from phasing.

Increasing the per homologue sequencing depth from 5-5-2× (mother-father-offspring) to 30- 30-30× decreased the SMR by 16-17% for SDhaP, PopPoly and TriPoly, while this decrease was 26% for H-PoP (Supplement S3).

#### PopPoly improves SNP dosage estimation

As shown in Figure 4, the incorrect dosage rate (IDR) in the phased SNPs was different for each method due to differences in each algorithm’s approach to handle genotype dosages.

**Figure 4.**
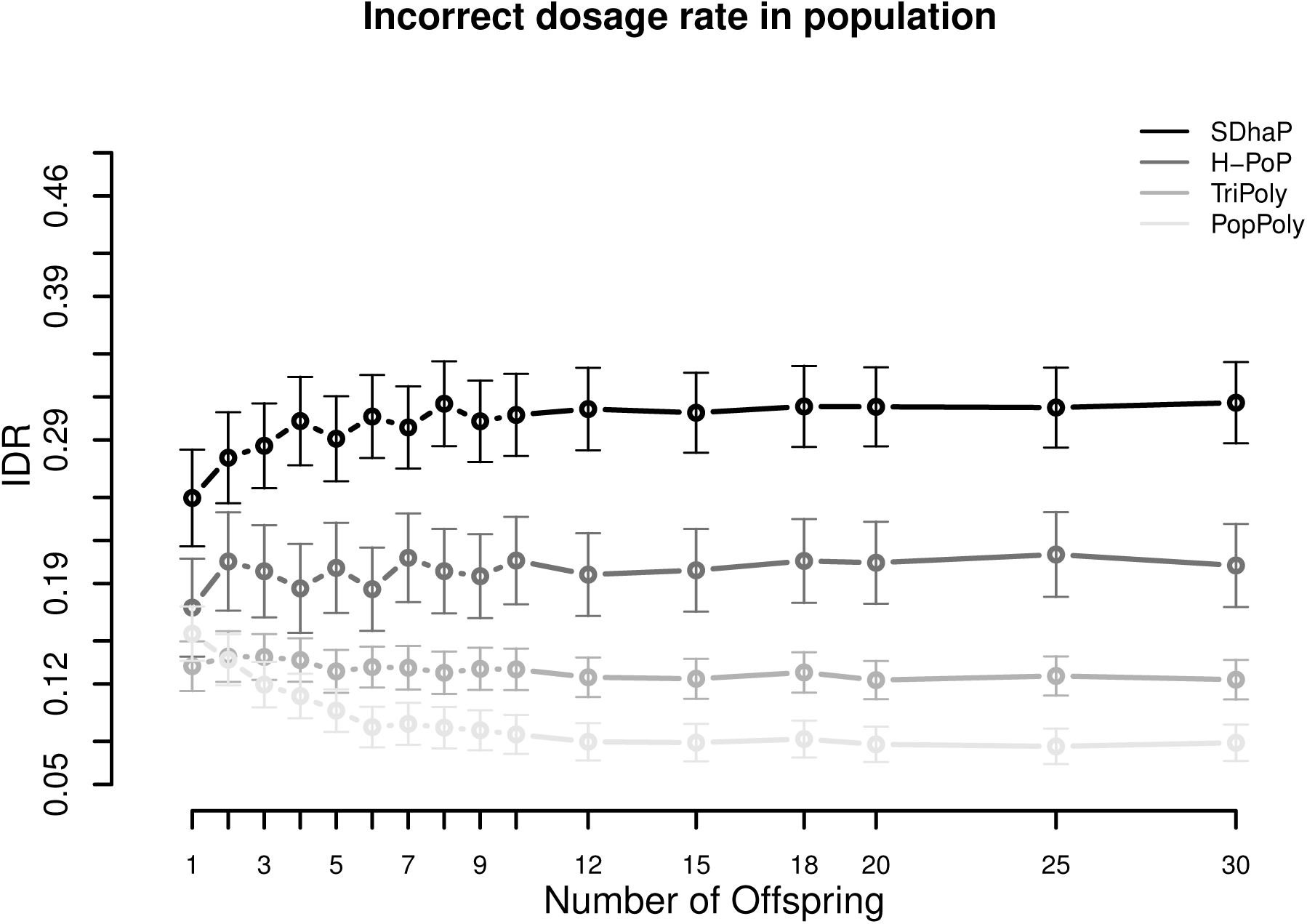
Incorrect dosage rate (IDR) in the population against the number of offspring reported by PopPoly (light grey), TriPoly (grey), H-PoP (dark grey) and SDhaP (black) for simulated tetraploid potato populations.

Specifically, H-PoP attempts to obtain an optimal partitioning of the reads into *k* groups corresponding to the homologues of a *k*-ploid, so that the difference between the reads assigned to the same homologue is minimised and the difference between the reads assigned to different homologues is maximised. The haplotypes are determined by taking a consensus of the reads within each group, and the dosages are determined by the estimated haplotypes. SDhaP on the other hand employs a gradient descent scheme with Lagrangian relaxation to find the best phasing (in the space of all possible phasings) according to the MEC criterion. Thus, its MEC solution determines the dosages of the SNP alleles.

In contrast to H-PoP and SDhaP, TriPoly and PopPoly use the input dosages as basis and make corrections to these based on parent-offspring relationships in the population. Specifically, if the genotype of an offspring in a trio is not compatible with the genotypes of the parents at position *s*, TriPoly obtains the offspring extension and hence the offspring genotype at *s* by considering all of the possible allele transmissions from the parents at *s* and by choosing the most likely trio extensions. The dosage correction method of PopPoly is explained in Section 2.2.

The simulation results show that the dosage correction scheme of PopPoly is the most successful approach if there are at least two offspring in the population (Figure 4). For a trio, however, the most accurate dosages are reported by TriPoly, while the IDR is the same for TriPoly and PopPoly with 2 offspring. On average, the IDR is around 31% for SDhaP (the highest), followed by 20% for H-PoP and 13% for TriPoly. With at least 6 offspring, the IDR of PopPoly drops below 10% (~7%).

As discussed for the phasing accuracy, the ability of PopPoly to detect wrongly estimated dosages and to correctly (re)estimate dosages depends on the haplotype counts in the population. Due to the absence of some parental haplotypes in the offspring of a trio, the accuracy of PopPoly drops below that of TriPoly, which relies less on the parental haplotypes and more on the read of the offspring to assign its dosages.

Considering the sequencing coverage, SDhaP profited the most from the higher depths with a 24% lower IDR at 30-30-30× compared to at 5-5-2× (per homologue), while this decrease in IDR was 12% for TriPoly and H-PoP and only 7% for PopPoly (Supplement S3).

#### Continuity of haplotyping is improved by PopPoly compared to single individual methods

The number of haplotype blocks, i.e. the number of gaps in an estimated phasing plus one [17], for a set of SNPs, 𝒮, is equal to the number of connected components in the *SNP-connectivity graph*, 𝒢_*S*_= (𝒮*, E_S_*). Each node in 𝒢_*S*_represents a SNP and an edge is drawn between two SNP nodes, (*s*, *s′*), if *s* and *s′* are covered together by at least one sequence fragment (which could be a single read or a paired-end sequence fragment). As shown in Figure 5, the expected number of phasing gaps (normalised by the number of SNPs, |𝒮|) is much lower in the estimates of TriPoly and PopPoly compared to H-PoP and SDhaP, as a pair of SNPs has a higher chance of being connected when all of the population reads are considered for the phasing of each individual compared to the case where for each individual only its own reads are taken into account.

**Figure 5.**
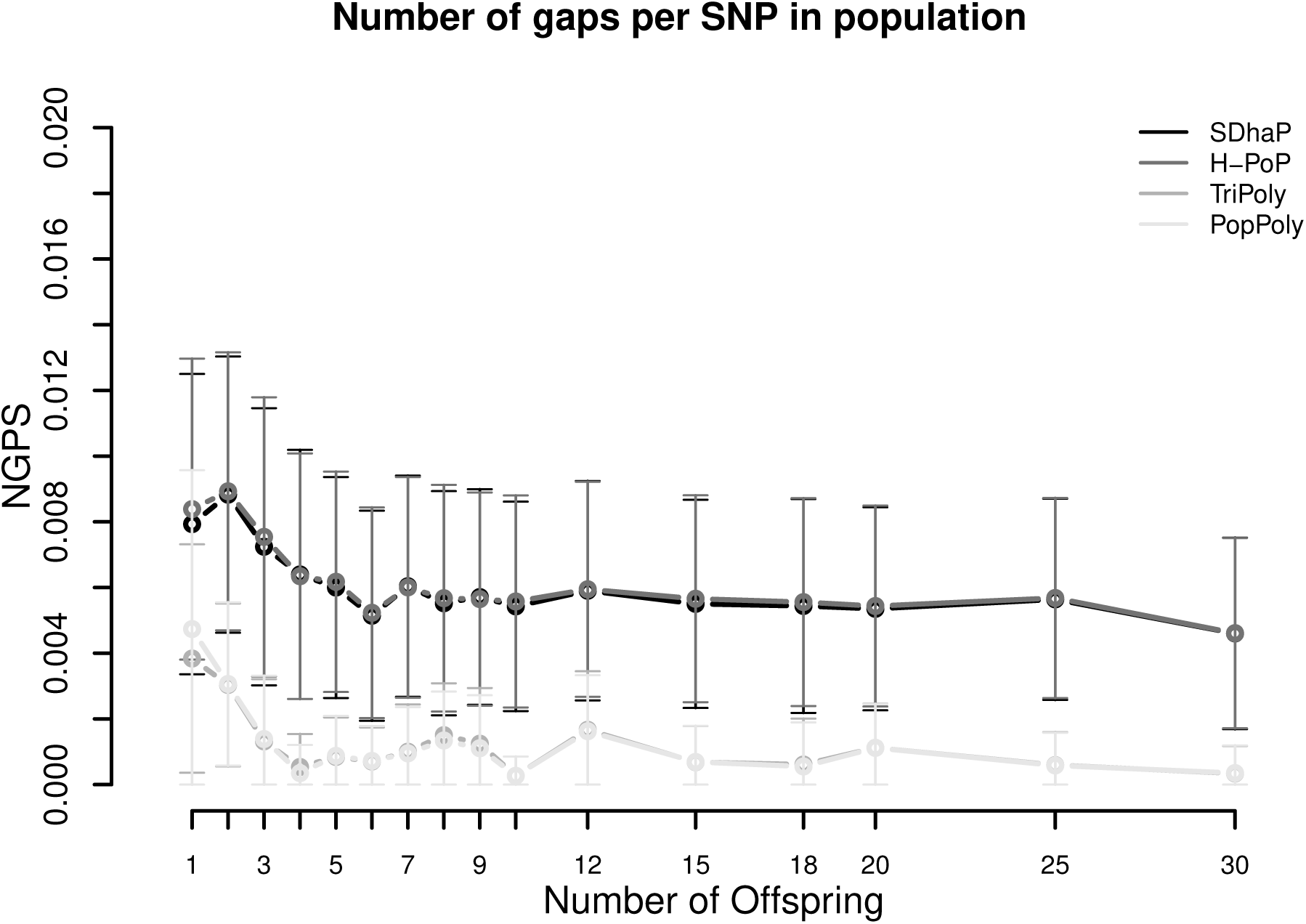
Number of phasing gaps normalised per SNP (NGPS) in the haplotype estimates of PopPoly (light grey), TriPoly (grey), H-PoP (dark grey) and SDhaP (black) against the number of offspring in the population for simulated tetraploid potato populations.

### 3.2. Haplotypes of tuberisation and maturity loci in *A* × *C* population

Using PopPoly, we phased all of the 579 segregating SNPs at the 9 candidate loci (Supplement S2). For each locus, we used the estimated haplotypes to calculate nucleotide diversity [30], i.e. the expected chance of a nucleotide difference per site between two randomly chosen haplotypes in the population. The estimated parental haplotypes showed high local similarity, although globally, i.e. for the entire locus, they were often distinct (Table 2).

**Table 2.**
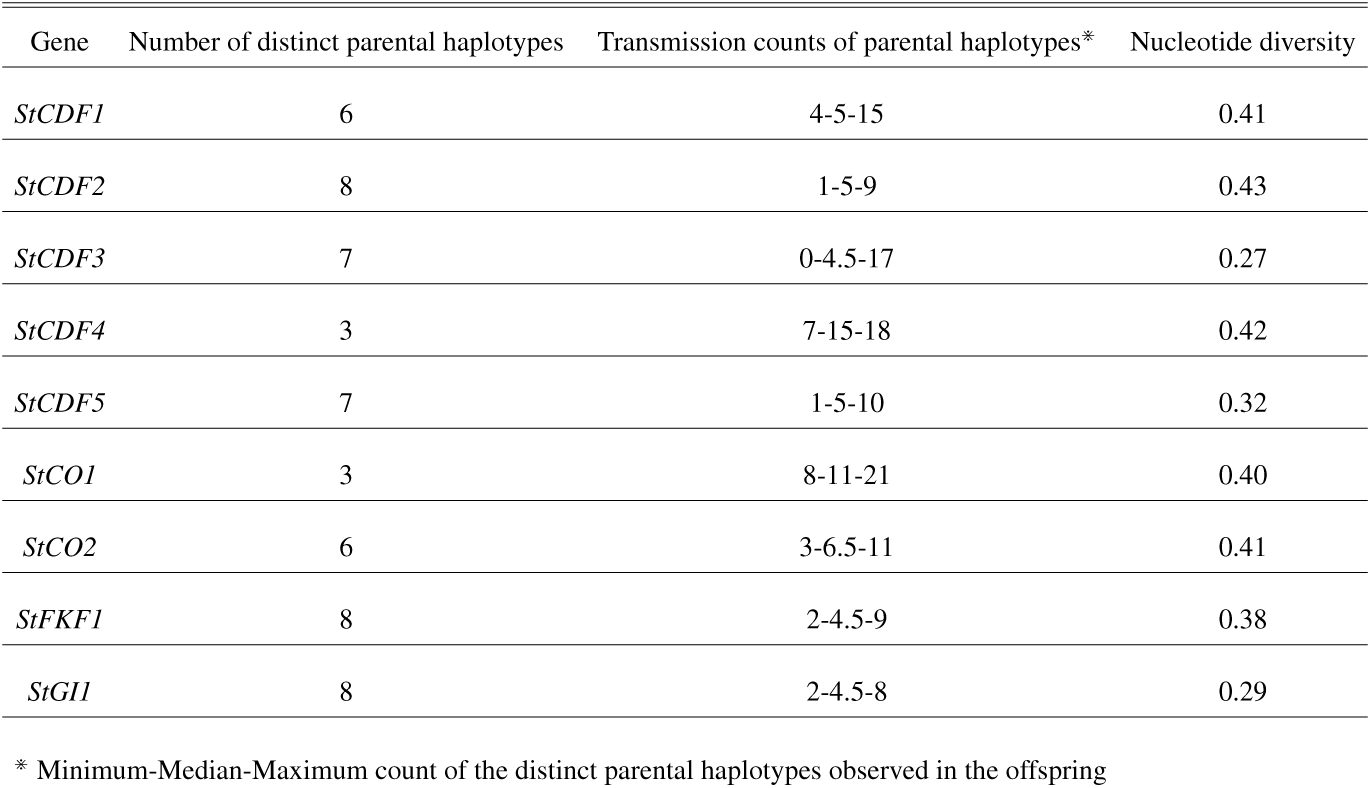
Summary of SNP phasing at the potato maturity and tuberisation loci (Table 1)

As evident from the median counts of the transmission of parental haplotypes to the offspring in Table 2, around half of the 56 distinct parental haplotypes (over all of the loci) were transmitted at least 5 times to the offspring. This is the expected transmission count of a haplotype in a tetraploid cross with 10 offspring if all of the parental haplotypes are distinct at the locus. However, one parental haplotype at *StCDF3* did not appear at all in the offspring estimates. A closer look at this locus (Table 3) shows that this haplotype, *H*_6_, is different from two other paternal haplotypes *H*_5_ and *H*_7_ (which are the same as each other) only at SNP sites *s* = 66 to *s* = 69, where *H*_6_ contains the reference alleles while *H*_5_ and *H*_7_ contain the alternative. With a larger population it will be possible to investigate whether this is due to phasing error or due to natural or human selection of the progeny. In comparison, the transmission pattern at *StCDF1* (Table 3) was as expected under the considered assumptions (Section 2.1).

**Table 3.** The 8 parental haplotypes and their transmission probabilities to each offspring at *StCDF1* and *StCDF3* loci (*H*_1_ – *H*_4_ represent maternal and *H*_5_ – *H*_8_ represent paternal haplotypes).

| Gene | Offspring id | $H_1$ | $H_2$ | $H_3$ | $H_4$ | $H_5$ | $H_6$ | $H_7$ | $H_8$ |
| --- | --- | --- | --- | --- | --- | --- | --- | --- | --- |
| *StCDF1* | 1 | 0 | 1 | 0 | 1 | 0.33 | 0.33 | 1 | 0.33 |
|  | 2 | 0 | 1 | 1 | 0 | 0.33 | 0.33 | 1 | 0.33 |
|  | 3 | 1 | 0 | 1 | 0 | 0.67 | 0.67 | 0 | 0.67 |
|  | 4 | 1 | 0 | 0 | 1 | 0.33 | 0.33 | 1 | 0.33 |
|  | 5 | 0 | 1 | 0 | 1 | 0.67 | 0.67 | 0 | 0.67 |
|  | 6 | 0 | 1 | 1 | 0 | 0.67 | 0.67 | 0 | 0.67 |
|  | 7 | 0 | 1 | 1 | 0 | 0.33 | 0.33 | 1 | 0.33 |
|  | 8 | 1 | 0 | 1 | 0 | 0.67 | 0.67 | 0 | 0.67 |
|  | 9 | 1 | 0 | 1 | 0 | 0.33 | 0.33 | 1 | 0.33 |
|  | 10 | 0 | 0 | 1 | 1 | 0.67 | 0.67 | 0 | 0.67 |
| *StCDF3* | 1 | 1 | 0 | 0 | 1 | 1 | 0 | 1 | 0 |
|  | 2 | 1 | 1 | 0 | 0 | 1 | 0 | 1 | 0 |
|  | 3 | 1 | 1 | 0 | 0 | 1 | 0 | 1 | 0 |
|  | 4 | 1 | 0 | 0 | 1 | 1 | 0 | 1 | 0 |
|  | 5 | 1 | 0 | 0 | 1 | 0.5 | 0 | 0.5 | 1 |
|  | 6 | 1 | 0 | 0 | 1 | 0.5 | 0 | 0.5 | 1 |
|  | 7 | 1 | 1 | 0 | 0 | 1 | 0 | 1 | 0 |
|  | 8 | 1 | 0 | 0 | 1 | 1 | 0 | 1 | 0 |
|  | 9 | 1 | 0 | 1 | 0 | 0.5 | 0 | 0.5 | 1 |
|  | 10 | 1 | 1 | 0 | 0 | 1 | 0 | 1 | 0 |

## 4. Conclusion

We present a novel algorithm, PopPoly, to exploit parent-offspring relationships for the estimation of haplotypes in F1-populations using short DNA sequence reads. Through realistic simulations, we show that PopPoly outperforms single individual haplotyping methods, which ignore family relationships. Besides, PopPoly yields better estimates compared to the trio based haplotyping method TriPoly when there are more than 2 offspring in the population. In addition, PopPoly employs the family information to improve variant dosage estimation in the population at the detected SNP sites. We also show that the performance of PopPoly is less influenced by sequencing depth compared to the other methods.

To demonstrate the utility of PopPoly, we used it to phase all of the 579 SNPs segregating at 9 plant maturity and tuberisation loci in an F1 population of tetraploid potato, the *A × C* cross, with 10 offspring.

## Acknowledgments

This work was funded by the graduate school Experimental Plant Sciences (EPS) of the Wageningen University and Research. We are also grateful for the support offered by the project initiative “Novel genetic and genomic tools for polyploid crops” (project number BO-26.03-009-004).

## Competing Interest

The authors declare that they have no competing interests.

